# The merits of sustaining pneumococcal vaccination after transitioning from Gavi support – a modelling and cost-effectiveness study for Kenya

**DOI:** 10.1101/369603

**Authors:** John Ojal, Ulla Griffiths, Laura L. Hammitt, Ifedayo Adetifa, Donald Akech, Collins Tabu, Anthony G. Scott, Stefan Flasche

## Abstract

**Introduction:** Many low income countries soon will need to consider whether to continue pneumococcal conjugate vaccine (PCV) use at full costs as they transition from Gavi support. Using Kenya as a case study we assessed the incremental cost-effectiveness of continuing PCV use.

**Methods:** We fitted a dynamic compartmental model of pneumococcal carriage to annual carriage prevalence surveys and invasive pneumococcal disease (IPD) incidence in Kilifi, Kenya, and predicted disease incidence and related mortality for either continuing PCV use beyond 2022, the start of Kenya’s transition from Gavi support, or its discontinuation. We calculated the costs per disability-adjusted-life-year (DALY) averted and associated prediction intervals (PI).

**Results:** We predicted that overall IPD incidence will increase by 93% (PI: 72% - 114%) from 8.5 in 2022 to 16.2 per 100,000 per year in 2032, if PCV use is discontinued. Continuing vaccination would prevent 15,355 (PI: 10,196–21,125) deaths and 112,050 (PI: 79,620– 130,981) disease cases during that time. Continuing PCV after 2022 will require an estimated additional US$15.6 million annually compared to discontinuing vaccination. The incremental cost per DALY averted of continuing PCV was predicted at $142 (PI: 85 - 252) in 2032.

**Conclusion:** Continuing PCV use is essential to sustain its health gains. Based on the Kenyan GDP per capita of $1445, and in comparison to other vaccines, continued PCV use at full costs is cost-effective. These arguments support an expansion of the vaccine budget, however, affordability may be a concern.

**Funding:** Funded by the Wellcome Trust.

## Introduction

The majority of African countries have introduced the pneumococcal conjugate vaccines (PCVs) in their childhood immunization programmes which has led to a substantial reduction in pneumococcal disease ^1,2^. In Kilifi, a coastal area in Kenya with enhanced surveillance for bacterial diseases, overall invasive pneumococcal disease (IPD) decreased by 68% in the post vaccination period (2012-2016) in children aged <5 years ^3^.

Although PCVs are among the most expensive vaccines available, most African countries were not concerned about affordability or cost-effectiveness when deciding to introduce PCV as Gavi, the Vaccine Alliance, took over the majority of vaccine costs. However, countries are expected to transition from Gavi support and subsequently take over the full costs once their 3-year-average Gross National Income per capita exceeds $1580. Currently three African countries (Angola, Congo Rep. and Nigeria) are in the accelerated transition phase ^4^ and six more (Ghana, Ivory Coast, Lesotho, Sudan, Kenya and Zambia) are expected to join within the next five years. With the increase in PCV costs upon transition countries will need to independently assess the cost-effectiveness and the affordability of sustaining PCV use.

Kenya introduced the 10 valent PCV (PCV10) in 2011 with Gavi’s support and has recently entered the preparatory transition phase, which will see their current contribution of $0.21 per dose increase by 15% annually. In 2022 Kenya will enter the accelerated transition phase that gradually increases their cost contribution to the full Gavi price of $3.05 by 2027 and thereby increasing PCV costs by 15 fold compared to current expenditure ^4^. Hence, before entering the accelerated transition-phase Kenya will need to evaluate whether to continue with PCV and or discontinue. We here assess the incremental impact and cost-effectiveness of continuing.

## Methods

We used a dynamic pneumococcal transmission model in combination with a costing model to estimate the cost-effectiveness of the two major policy options for PCV use in Kenya from 2022; i.e. continuation of PCV use at Gavi’s current and scheduled prices or discontinuing the vaccine. The approach accounts for the uncertainty in both epidemiology and costing estimates and propagates it to the predicted outcomes.

### Disease model and incidence prediction

The details of the transmission model have been described elsewhere ^5^. In brief, we used a compartmental, age-structured, dynamic model (Appendix, Supplementary Figure 3). The model has a Susceptible-Infected-Susceptible (SIS) structure for three serotype groups: the vaccine serotypes (VT), strongly competitive non-vaccine serotypes (sNVT) and weakly competitive non-vaccine serotypes (wNVT). We calibrated the model to age-stratified annual pre-vaccination (2009-2010) and post-vaccination (2011-2016) pneumococcal carriage prevalence by fitting serotype competition, susceptibility to infection if exposed and vaccine efficacy using non-informative priors for all parameters except the vaccine efficacy (Appendix).

In Kilifi, PCV vaccination was introduced together with a catch-up campaign in children <5 years old. To extrapolate findings to the rest of Kenya, where PCV was introduced without a catch-up campaign, the fitted model was re-run under these conditions. We predicted carriage incidence for a 15-year period, from 2017 to 2032. We predicted IPD incidence by multiplying modelled age-specific carriage incidence with case-to-carrier ratios (CCR). For each model posterior the CCRs were calculated as the ratio of the observed pre-vaccination IPD incidence at Kilifi Country Hospital (KCH)^3^ to modelled pre-vaccination carriage incidence. The CCR were assumed to remain unchanged post-vaccination.

IPD was defined as isolation of *Streptococcus pneumoniae* from a sterile site culture in an individual admitted to KCH. We split the predicted IPD incidence into the age dependent proportions that are pneumococcal meningitis, pneumococcal sepsis and bacteraemic pneumococcal pneumonia incidence based on the distribution observed in clinical data from KCH (Supplementary Table S1). We defined pneumococcal meningitis as isolation of *Streptococcus pneumoniae* from cerebrospinal fluid (CSF) or isolation of *S. pneumoniae* from blood, accompanied by a CSF white blood cell count of 50 x 10^6^ cells/L or greater or a ratio of CSF glucose to plasma glucose less than 0.1. Bacteraemic pneumococcal pneumonia was defined as IPD with no pneumococcal meningitis but with WHO severe or very severe pneumonia. Pneumococcal sepsis was defined as IPD not meeting the definitions of pneumococcal meningitis or bacteraemic pneumococcal pneumonia. We further assumed that for every prevented case of IPD one would prevent 5.3 cases of clinically-defined pneumonia ^3,6^. This ratio was estimated by dividing the vaccine preventable clinical pneumonia incidence (351 per 100,000 per year) ^3^ to vaccine preventable IPD incidence (66.3 per 100,000 per year) ^6^ that were both estimated from surveillance at KCH. The hospital surveillance in KCH was found to underestimate the incidence of pneumonia and meningitis by 45% and 30% respectively ^7^. We accounted for this age-independent under reporting in our analysis by inflating case numbers commensurately.

### Vaccination program costs

The program costs included vaccine costs, vaccine wastage, safety boxes, administering syringes for each dose, reconstitution syringes for each vial, syringe wastage and vaccine delivery cost (Table 1). Vaccine cost used for each year was calculated according to Gavi transitions rules (Supplementary Table 2). The vaccine delivery cost included the vaccine supply chain cost and immunization service delivery cost. The initial investment in expanding the cold chain capacity in 2011 was not included. A switch from 2-dose to 4-dose presentation occurred in 2017. The 4-dose presentation has a preservative and once opened for the first time the vial can be kept for up to 28 days, therefore, no noteworthy change in vaccine wastage rates is expected ^8^.

**Table 1:** Economic and health parameters varied as part of the probabilistic sensitivity analysis

| Parameter | Point estimate | Statistical distribution | Source |
| --- | --- | --- | --- |
| **Access to care proportions for pneumococcal diseases** |  |  |  |
| Hospitalised sepsis, bacteraemic and non-bacteraemic cases | 55% | Beta (55,45) | <sup>7</sup> |
| Hospitalized meningitis cases | 70% | Beta (70,30) | <sup>7</sup> |
| Non-hospitalised IPD and non-bacteraemic cases treated as outpatient | 63% | Beta (63,37) | <sup>20</sup> |
| **Health outcomes** |  |  |  |
| Proportion of meningitis cases developing sequelae | 25% | Beta (25,75) | <sup>21</sup> |
| CFR with hospital care |  |  |  |
| Sepsis, bacteraemic and meningitis: Children (<15 years) | 19% | Beta (19,81) | KCH |
| Sepsis, bacteraemic and meningitis: Adults (≥15 years) | 46% | Beta (46,54) | KCH |
| Non-bacteraemic pneumonia | 5.7% | Beta (6,94) | <sup>22</sup> |
| CFR without hospital care |  |  |  |
| Meningitis | 97% | Beta (97,3) | <sup>23</sup> |
| Sepsis and bacteraemic pneumonia | 50% | Beta (4,4) | <sup>23</sup> |
| Non-bacteraemic pneumonia | 12% | Beta (12,88) | <sup>23</sup> |
| **Vaccination costs (US\$)** | | | |
| Vaccine price per dose | \$0.21-\$3.05 (Table S2) | Fixed | <sup>4,24,25</sup> |
| Safety boxes | \$0.46 | Fixed | <sup>26</sup> |
| AD syringes | \$0.045 | Fixed | <sup>26</sup> |
| Vaccine delivery cost per dose | \$1.42 | Gamma (4,0.4) | <sup>27</sup> |
| Syringe wastage | 5% | Fixed | <sup>23</sup> |
| Vaccine wastage | 15% | Fixed | <sup>8,23,28</sup> |
| **Treatment costs (US\$)** | | | |
| With hospital care |  |  |  |
| Meningitis | \$357.74 | Gamma (4,97) | <sup>29</sup> |
| Sepsis, bacteraemic and non-bacteraemic pneumonia | \$74.64 | Gamma (4,19) | <sup>29</sup> |
| With outpatient care (All four syndomes) | \$2.74 | Gamma(4,0.75) | <sup>30</sup> |
| Without hospital care (All four syndomes) | \$1.15 | Gamma (4,0.3) | <sup>30</sup> |

### Treatment costs

We adopted a societal perspective in our analyses, i.e. including direct medical costs, the opportunity cost of caretaker time and household out-of-pocket costs.

To apply the appropriate treatment costs, we divided the cases into three groups depending on where they were treated: hospitalised cases, cases treated as outpatients and those that did not reach medical care (Table 1). All costs not referring to 2016 were converted into 2016 US dollars for our analysis by using the International Monetary Fund’s (IMF) GDP deflators for Kenya.

### Disability Adjusted Life Years (DALYs)

The treatment costs for the predicted number of cases for the four syndromes considered and the vaccination cost of birth cohorts were estimated and used to calculate the costs per disability-adjusted-life-year (DALY) averted. The years lost due to disability (YLD) were calculated as the product of disease incidence, duration of disease and disability weights. We used disability weights from the 2013 global burden of disease study ^9^ in calculating YLD component of DALYs. We used the disability weight of 0.133, assigned for infectious diseases with severe acute episodes, for both IPD and non-bacteraemic pneumonia episodes. For meningitis sequelae, we used a disability weight of 0.542 assigned for motor plus cognitive impairment. We assumed a duration of 15 days for all IPD syndromes and 7 days for non-bacteraemic pneumonia. Meningitis sequelae were assumed to last a lifetime. We used the Kenyan age specific life expectancies ^10^ in calculating the Year of Life Lost (YLL) due to death. The discount rate on costs and DALYs was set at 3%.

### Sensitivity analysis of the cost inputs and disease model

The full uncertainty of both epidemiological and costs parameters was propagated to the results as follows: for each posterior estimate of the epidemiological model we sampled a set of cost parameters from the pre-set distributions, effectively combining probabilistic fitting of the epidemiological mode with a probabilistic sensitivity analysis of the costing model (Table 1).

In Kenya, children who are carriers of VT pneumococci have been observed to respond less well to vaccine than non-carriers ^11^. To assess structural uncertainty in our model we ran our analyses either with or without accounting for hyporesponsiveness. In the base case, we estimated a single vaccine efficacy independent of carrier status; in the sensitivity analysis, vaccine efficacy was estimated separately in vaccine-type carriers and in others. We also present two scenarios of discounting, i.e. discounting both costs and DALYs at 3% (base case) or discounting costs alone.

## Results

### Model fit and predicted IPD incidence

There was good agreement between the observed and fitted age-group and serotype-group specific carriage prevalence (Figure 1 & Appendix) and IPD incidence (Figure 2). If cohorts of children born after the start of year 2022 are no longer vaccinated with PCV, the model predicts that IPD incidence will bounce back from 8.5, in 2022 to 16.2 per 100,000 per year in 2032 equalling pre PCV levels (Figure 3). Continuing with PCV is predicted to result in additional small reductions in IPD incidence to 7.9 per 100,000 per year in 2032, and to avert 15,355 (PI: 10,196–21,125) deaths and 112,050 (PI: 79,620–130,981) IPD and non-bacteraemic pneumonia cases during the 11 years considered, compared to discontinuing the PCV in 2022.

**Figure 1:**
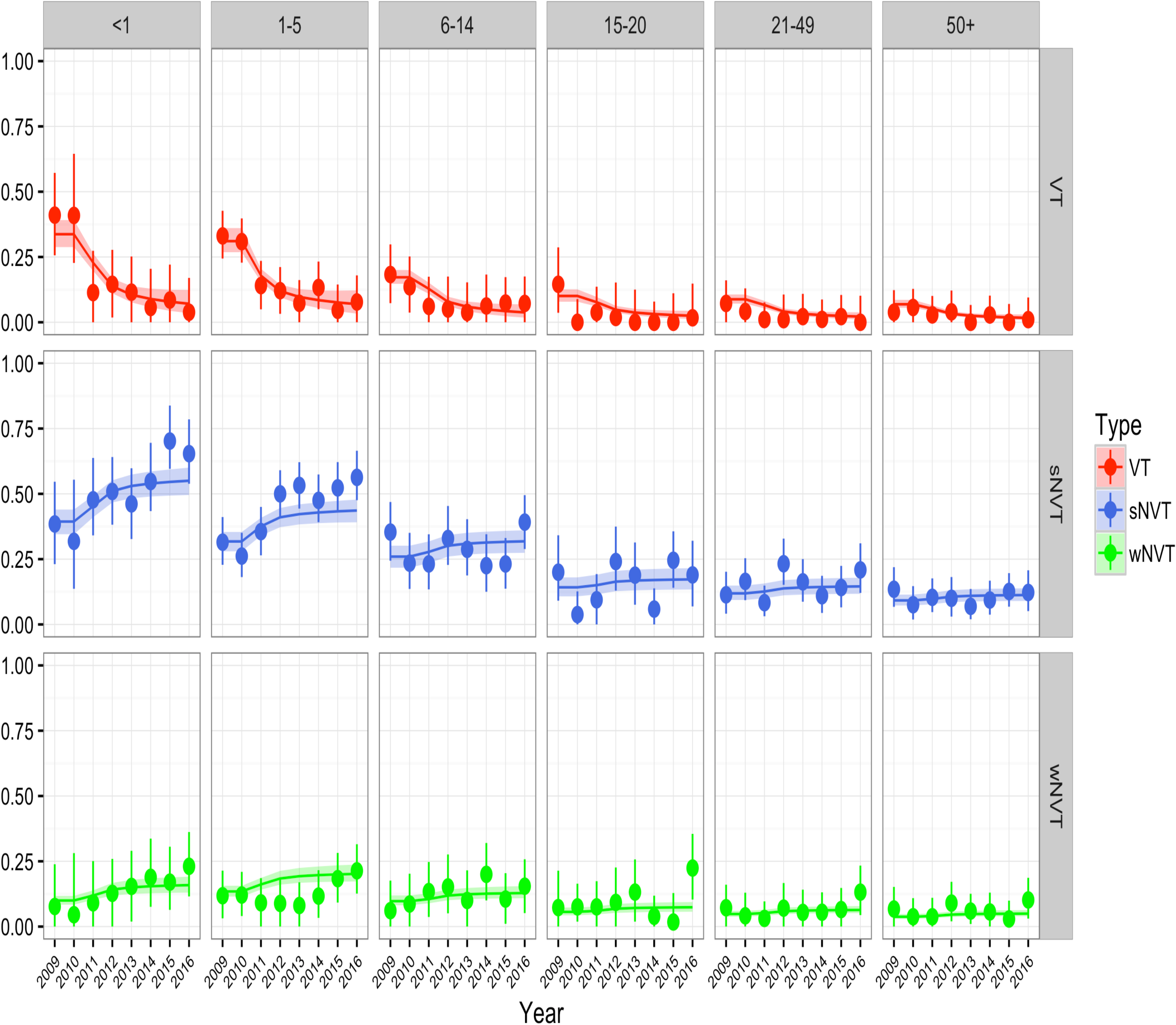
Model fit to carriage data. Observed (circular dots with 95% credible intervals shown by spikes) and predicted (lines with 95% predictive intervals shown by shaded areas) carriage prevalence of vaccine-serotypes (VT), shown in red, strong non-vaccine serotypes (sNVT), shown in blue, and weak non-vaccine serotypes (wNVT), shown in lime green, over time. The age groups are labelled at the panel titles.

**Figure 2:**
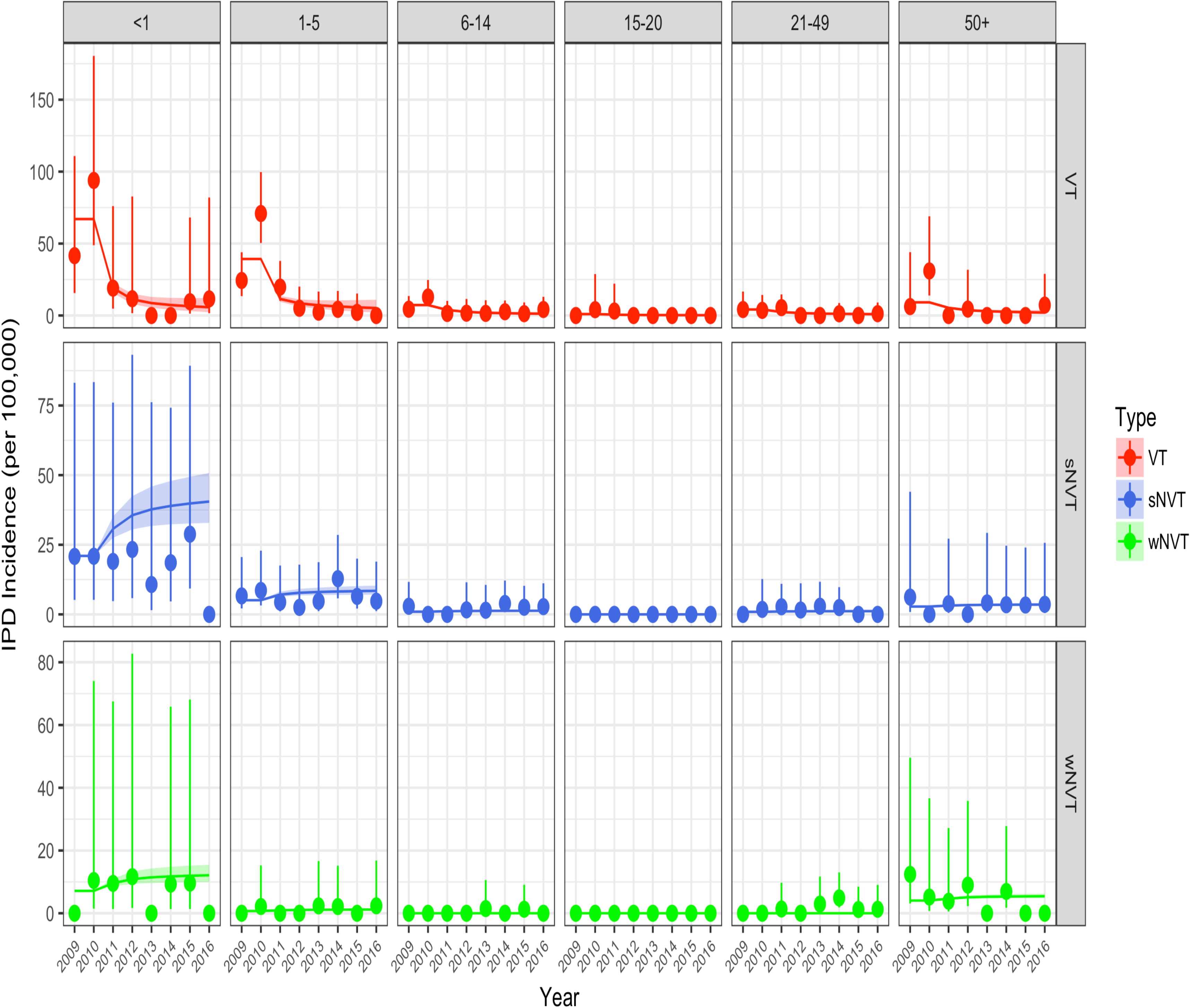
Model fit to IPD incidence data. Observed (circular dots with 95% credible intervals shown by spikes) and predicted (lines with 95% predictive intervals shown by shaded areas) IPD incidence of vaccine-serotypes (VT), shown in red, strong non-vaccine serotypes (sNVT), shown in blue, and weak non-vaccine serotypes (wNVT), shown in lime green, over time. The age groups are labelled at the panel titles.

**Figure 3:**
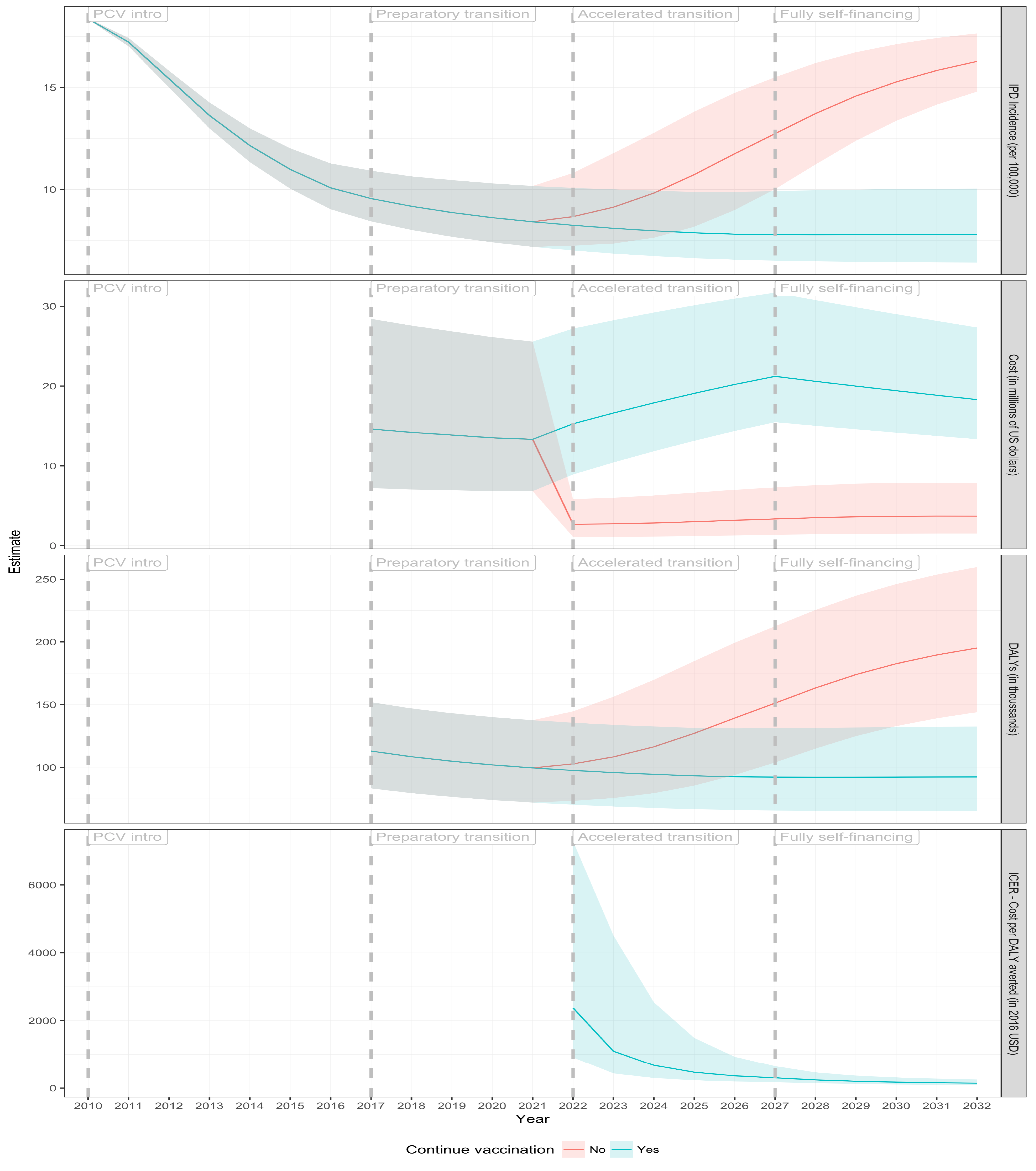
Costs, DALYs, incremental cost-effectiveness ratios (ICERs) and invasive pneumococcal disease (IPD) incidence at the end of each year. The first (top-most) panel shows the predicted incidence of IPD when vaccination is continued in 2022 (cyan line), its 95% prediction interval (cyan shade), and when vaccination is stopped (red line, with 95% prediction interval shown in light-red shade) over time since vaccine introduction in Kenya at the end of 2010. The second panel shows the cost of treatment and vaccination in each year (cyan line with cyan shade for 95% PI) when vaccination is continued, and the cost of treatment in each year when vaccination (red line, with light-red shade for 95% PI) is discontinued in 2022. The third panel shows the corresponding DALYs gained in each year. The fourth (bottom-most) panel shows the ICER (y-axis), incremental (continuing vaccination over stopping vaccination) cost per DALY averted (cyan line) and its 95% prediction interval (cyan shade) in each year (x-axis). Vertical dotted lines indicate Gavi transition stages.

### Estimated costs and cost effectiveness

If vaccination was to be stopped in 2022 the estimated average annual treatment cost for pneumococcal disease in Kenya would be $3,275,143. Otherwise, average annual treatment and vaccination costs for continuing PCV during 2022-2032 were estimated as $18,851,991 (Table 2). Discontinuing PCV was predicted to partially sustain direct and indirect protection from the vaccination of previous cohorts for some of the study period with gradually declining impact on IPD incidence. As a result, we predict that continuation of PCV will not be cost effective initially. However, we show that within only one year after the decision to continue PCV the incremental cost-effectiveness ratio (ICER), in comparison to discontinuing PCV, improves substantially towards the threshold of the Kenyan GDP per capita ($1455 in 2016) and continues to improve throughout the study period (Figure 3). Compared to discontinuing PCV in 2022, we predicted that, in 2032, the cost per DALY averted is $142, the cost per case averted $876 and the cost per death averted $6366 (Table 2).

**Table 2:** Estimated costs and cost-effectiveness ratios for different scenarios

| Scenario | Average annual cost<br>Over 2022-2032, millions of US\$<br>(95% PI) | Cost per case<br>averted in 2032, US\$<br>(95% PI) | Cost per death<br>averted in 2032, US\$<br>(95% PI) | Cost per DALY<br>Averted in 2032, US\$<br>(95% PI) |
| --- | --- | --- | --- | --- |
| Stopping vaccination in year 2022 | 3.3 (1.3 – 7.1) | Ref. | Ref. | Ref. |
| Continuing vaccination | 18.9 (13.2 – 29.3) | 876 (564- 1443) | 6366 (3802 – 11226) | 142 (85 – 252) |
| Continuing vaccination (discounting costs only) | 18.9 (13.2 – 29.3) | 529 (341 – 873) | 3852 (2300 – 6792) | 67 (40 – 120) |

### Sensitivity analyses

Using the Kenyan GDP per capita of $1455 in 2016 as a threshold to determine cost effectiveness, all posterior samples indicated that continuation of PCV vaccination is cost effective no more than six years after 2022. Compared to discounting both costs and DALYs, discounting costs alone resulted in an ICER that was twice as favourable (Table 2).

We estimate that the effect of hyporesponsiveness is relatively small. Vaccine serotype carriers had a vaccine efficacy estimate against carriage that was 4 percentage points lower than that for other vaccinees (Appendix, table A1). Hence omitting this mechanism in the model structure led to similar results (Supplementary Figure 3). Therefore, we did not include hyporesponsiveness in our final model.

## Discussion

In the near future Kenya, like several other low income countries, will be expected to take over the full cost of the national pneumococcal conjugate vaccination procurement. In this study, we have estimated the cost-effectiveness of continuing PCV using Gavi’s schedule of vaccine prices, which reach a peak at $3.05 per dose in 2027, at which point Kenya becomes fully self-financing. Our model projects that discontinuing PCV would lead to an increase in IPD burden equivalent to pre-vaccination levels within ten years. Initially, continuing vaccination may not be cost-effective because of the benefits accrued through vaccination of previous cohorts. However, the cost-effectiveness becomes substantially more favourable within a few years and, by 2032, the cost (in 2016 US dollars) plateaus at $142 ($85 -$252) per discounted DALY averted.

The most commonly used threshold for judging the cost-effectiveness of an intervention is a country’s Gross Domestic Product (GDP) per capita. Using this criterion, we find continuation of PCV in Kenya after transition from Gavi support highly cost-effective. The GDP per capita threshold was initially supported by the Commission on Macroeconomics and Health ^12^ and adopted by WHO’s Choosing Interventions that are Cost-Effective project (WHO-CHOICE). The use of GDP-based thresholds has been criticized because it: (i) does not consider the cost-benefits profile of interventions competing for the same health budget; (ii) does not adequately address the willingness to pay; (iii) does not address affordability and (iv) is easily attained. Alternatives include benchmarking of interventions by assessing a country’s willingness to pay by comparing cost-effectiveness ratios against that of vaccines currently in use.

The cumulative costs per DALY averted of introducing the Rotarix or the RotaTeq rotavirus vaccines in Kenya have been estimated as $200 and $406 (2016 US Dollars) respectively. Similar to our estimates these were derived based on a societal perspective with a 3% discounting of both costs and benefits ^13^. The *Haemophilus influenzae* type B (Hib) vaccine was introduced in 2001 Kenya as part of the pentavalent vaccine. In a static model developed to follow the Kenyan 2004 birth cohort until death, with and without Hib vaccine, it was estimated that the discounted (3% for both costs and benefits) cost per DALY averted of introducing Hib vaccine was $85 (2016 US Dollars) from a health provider perspective ^14^. This suggests that continuation of PCV is less cost-effective than the Hib vaccine and more cost-effective than the rotavirus vaccine. However, these comparisons must be tempered by the fact that the rotavirus analysis ignored herd immunity, while the Hib analysis took a health provider perspective, both of which decrease cost-effectiveness.

Cost-effectiveness, however, does not necessarily imply affordability. The later depends on available resources in the health budget, or any other sources within the national accounts that can fill the gap in the health budget. Budgetary allocation to the health sector as a fraction of national government budget has slightly declined from 4% in financial year 2014/15 to 3.7% in financial year 2016/17 ^15^. The Kenyan annual health budget for 2015 was $600 million ^15^. Out of this $6.9 million (0.8%)^16^ was spent on vaccines. This has been possible because Kenya only needs to fund 10% of its vaccines from its revenues, donors fund the rest of the budget ^17^. We have estimated that continuing with PCV after 2022 will require an additional $15.6 million annually compared to discontinuing PCV; in other words, it will more than double Kenya current expenditure on vaccines. At the same time, following transition from Gavi support, the Kenyan government financial contribution for pentavalent, rotavirus and yellow fever vaccines, will need to increase as well if Kenya wants to sustain their current vaccine portfolio, putting further stress to the budget.

Several initiatives indicate that the cost of the PCV procurement may be reduced in future. For instance, the Serum Institute of India is developing of a 10-valent PCV with a target per-dose price of $2.00 ^18^. Also, in settings where vaccine serotypes have been eliminated from circulation it may be possible to sustain control of transmission using a two-dose or even one-dose schedule^19^. If vaccine serotypes can be eliminated in Kenya, for example by additional efforts such as a catch-up campaign, then the shift to a reduced dose schedule may also be feasible. Most of these options will have a wider evidence base that may allow their formal consideration by 2022. Currently there is insufficient support to include them in our analyses but if proven to be effective these aspects will further improve on our PCV cost-effectiveness estimates of sustaining PCV in Kenya.

There are potential limitations to our study. The proportion of pneumococcal disease cases that are hospitalized, treated as outpatients or do not access care is a key determinant of both costs incurred as well as DALYs, by determining the case fatality rate. Overestimating the proportion of cases that get hospital treatment would mean that the overall costs of treatment were overestimated while the fatal cases, and therefore DALYs, were underestimated. The overall effect would be an overestimated ICER, which is conservative. In our analysis, we estimated the proportion of cases that were hospitalized using local surveillance data. However, we did not have local information on what proportion among non-hospitalised cases are treated as outpatients; this was obtained from a Ugandan verbal autopsy study among fatal pneumonia cases ^20^. It is possible, therefore, that we have overestimated the number of patients among non-hospitalised treated as outpatients, and, by extension, overestimated the ICER.

Several low-income countries will soon be transitioning out of Gavi support and will need to decide whether to sustain their pneumococcal conjugate vaccination. We demonstrate, using Kenya as an example, how ongoing detailed surveillance can be combined with mathematical modelling and health economics to inform an upcoming decision of a country’s National Immunization Technical Advisory Group (NITAG) on the cost-effectiveness of different policy options. We estimate that maintaining PCV is essential to sustain the decreased burden of pneumococcal disease and that it is cost-effective against conventional criteria. However, to afford PCV vaccination in the post-Gavi era, Kenya will need to substantially increase the proportion of health spending on routine immunization.

## Author Contributions

The study was conceived by UG and JAGS. The model was designed by JO and SF and coding and simulations were by JO. LLH, DA, IA, JAGS conducted the pneumococcal carriage surveys and/or oversaw the IPD surveillance. JO wrote the first draft of the manuscript. All authors read and critically reviewed the manuscript and approved the final version.

## Ethics statement

The study was part of the Pneumococcal Conjugate Vaccine Impact Study (PCVIS) approved by the Kenya Medical Research Institute (KEMRI) Ethical review committee (SSC 1433). It has an additional approval by OXTREC (OXTREX 30-10), the Oxford Tropical Research Ethics Committee, with delegated authority from the London School of Hygiene & Tropical Medicine (LSHTM) Research Ethics Committee.

## Acknowledgements

This paper is published with the permission of the Director, KEMRI. The work was funded by the Wellcome Trust through fellowship support to John Ojal (092767) and Anthony Scott (098532). Stefan Flasche is supported through a Sir Henry Dale Fellowship jointly funded by the Wellcome Trust and the Royal Society (Grant Number 208812).

---

**Supplementary table S1:** Invasive Pneumococcal disease (IPD) separation into meningitis, pneumococcal bacteraemic pneumonia and sepsis. IPD cases are obtained from hospitalized cases at the Kilifi County hospital among residents of the Kilifi Health and Demographic Surveillance System for the period 1999-2016 (<15) for children and 2007- 2016 for adults (>=15).

| Age category | Pneumococcal Meningitis<sup>a</sup> |  | Pneumococcal Bacteraemic Pneumonia<sup>b</sup> |  | Pneumococcal Sepsis<sup>c</sup> |  |
| --- | --- | --- | --- | --- | --- | --- |
|  | n | % | n | % | n | % |
| <1 | 52 | 28.0 | 85 | 45.7 | 49 | 26.3 |
| 1-5 | 32 | 11.3 | 126 | 44.4 | 126 | 44.4 |
| 6-14 | 39 | 29.8 | 47 | 35.9 | 45 | 34.4 |
| 15-20 | 1 | 33.3 | 0 | 0.0 | 2 | 66.7 |
| 21-49 | 11 | 29.7 | 3 | 8.1 | 23 | 62.2 |
| 50+ | 3 | 11.5 | 2 | 7.7 | 21 | 80.8 |

<sup>a</sup> Isolation of *S. pneumoniae* from cerebrospinal fluid (CSF) or isolation of *S. pneumoniae* from blood, accompanied by a CSF white blood cell count of $50 \times 10^6$ cells/L or greater or a ratio of CSF glucose to plasma glucose less than 0.1
<sup>b</sup> IPD with no pneumococcal meningitis but with WHO severe or very severe pneumonia.
<sup>c</sup> IPD not meeting any of the above definitions of pneumococcal meningitis and bacteraemic pneumococcal pneumonia.

**Supplementary table S2:**
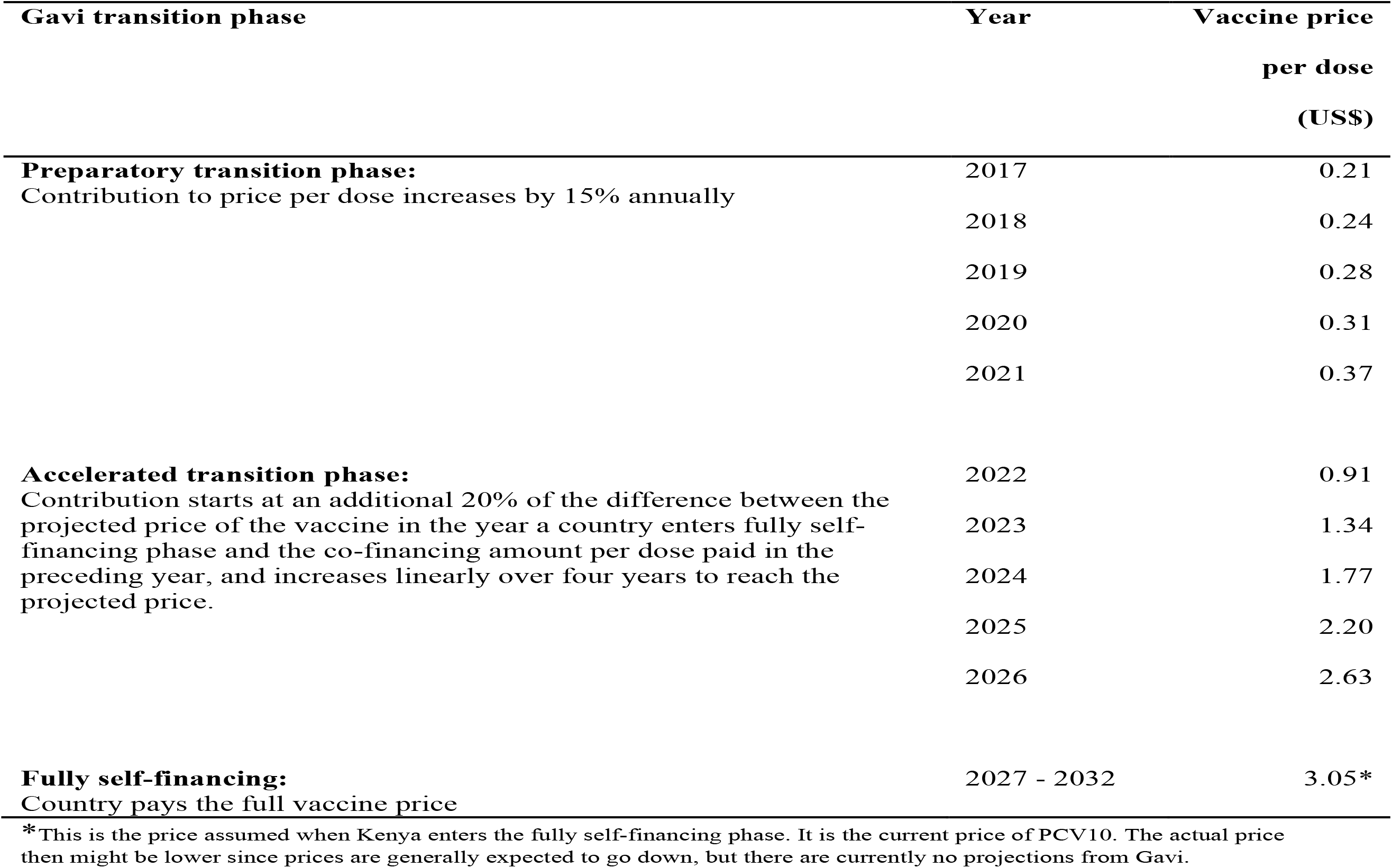
Vaccine price per dose paid by Kenya in each year.

**Supplementary Figure 1:**
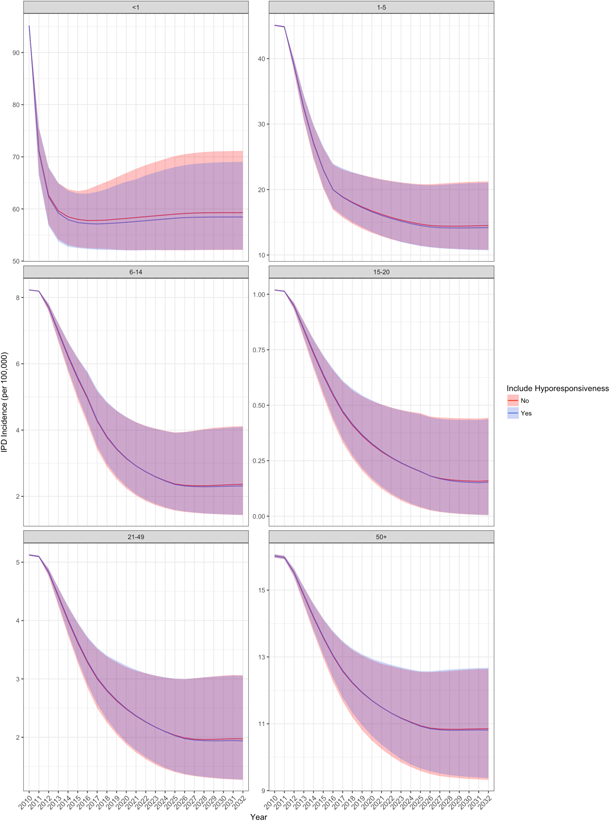
IPD projection with and without hyporesponsiveness by age group. Predicted incidence of IPD when hyporesponsiveness is ignored in the carriage model (red line with 95% prediction interval shown in light-red shade, and when hyporesponsiveness is allowed for in the model structure (blue line, with 95% prediction interval shown in light-blue shade) over time since vaccine introduction in Kenya. Age groups are labeled on the panel titles.

**Supplementary Figure 2:**
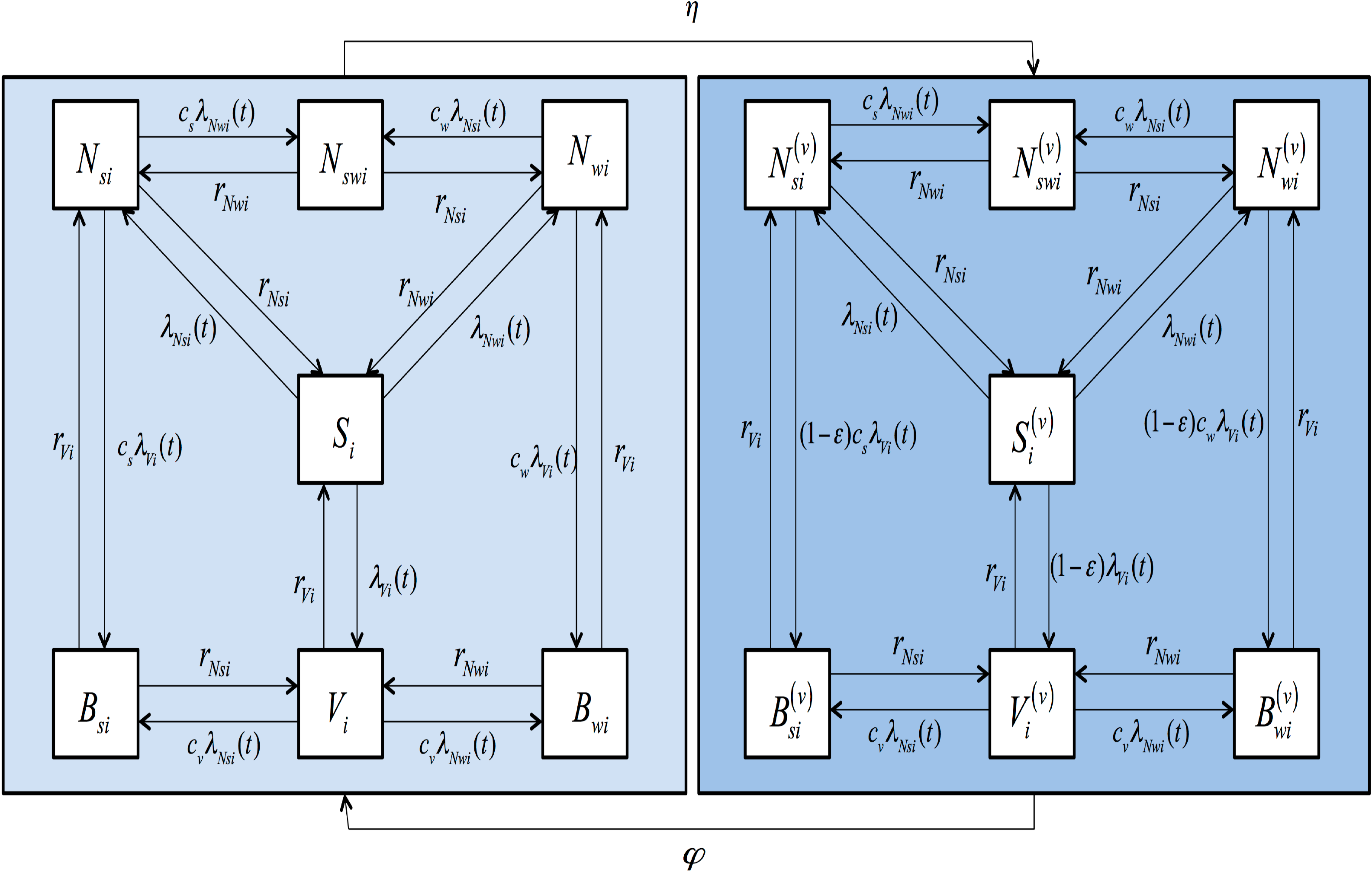
Model structure flow diagram. The epidemiological states include individuals that are susceptible (non-carrying), *S*; carry a vaccine serotype, *V*; carry a weak non-vaccine serotype, *N_W_*; carry a strong non-vaccine serotype, *N_S_*; carry simultaneously a weak and a strong non-vaccine serotype, *N_SW_*; carry simultaneously a vaccine serotype and a weak non-vaccine serotype, *B_W_*; or carry simultaneously a vaccine serotype and a strong non-vaccine serotype, *B_S_* (see text). Once vaccinated, the individual moves to one of the corresponding states, (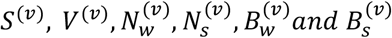). The acquisition rates from the single to multiple serotype carriage states are reduced by competition parameters denoted by *c* with two subscripts; the first denoting the serotype group (*υ*, *s* and *w* for VT, strong NVT and weak NVT respectively) of the resident serotypes and the second denoting the age-group. The competition parameters have two sets of values, one for age group <6 and another for age group ≥6 years (see Appendix). The age-group specific VT, weak NVT and strong NVT clearance rates are denoted by r_Vi_, r_Nwi_ and r_Nsi_, respectively. In addition to the transitions between the 14 epidemiological states as shown in the Figure, individuals die from any states at age-specific death rates and new individuals are born into the completely susceptible state.

**Supplementary Figure 3:**
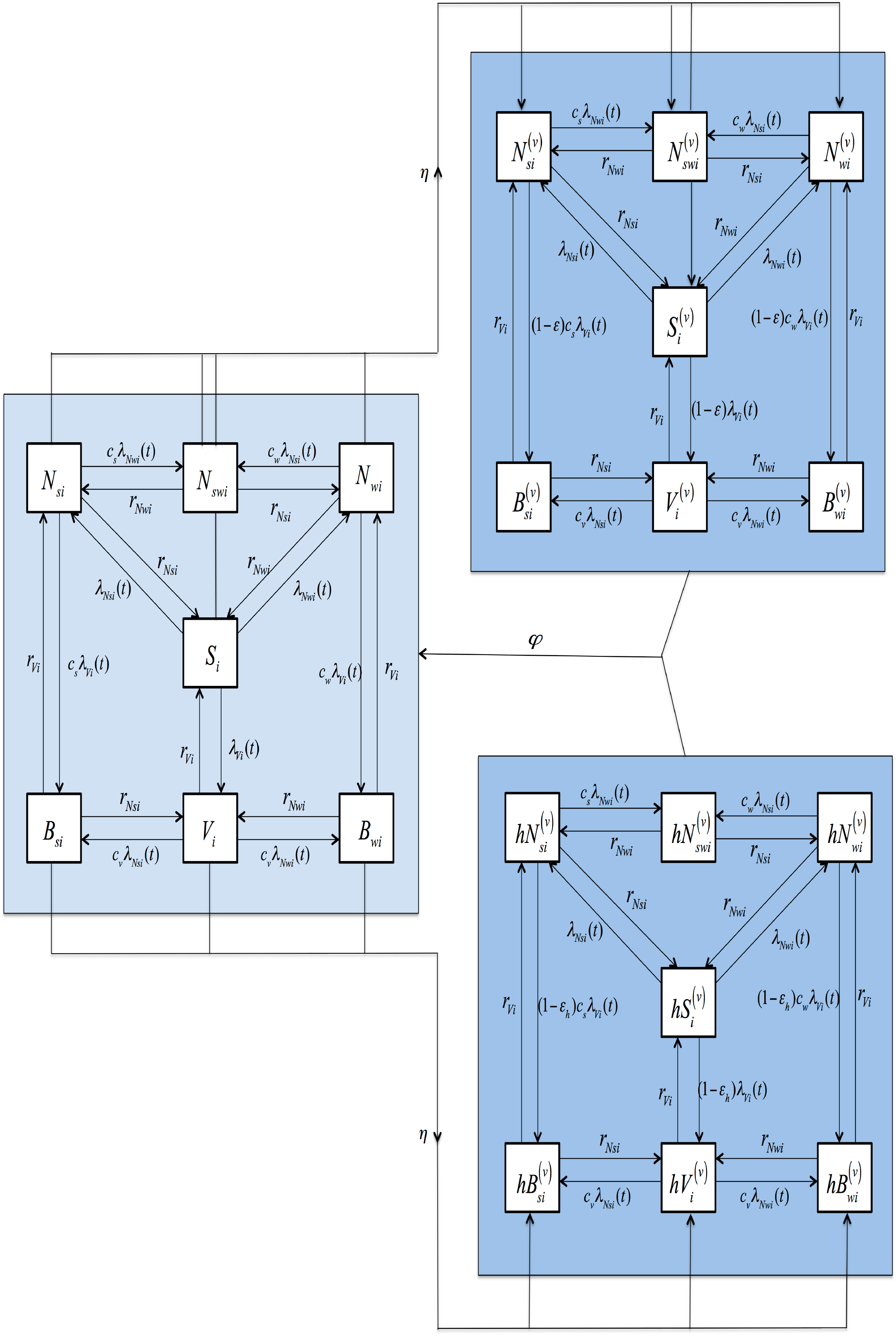
Model structure flow diagram including hyporesponsiveness. The epidemiological states include individuals that are susceptible (non-carrying),*S*; carry a vaccine serotype, *V*; carry a weak non-vaccine serotype, *V*; carry a strong non-vaccine serotype, *N_w_*; carry simultaneously a weak and a strong non-vaccine serotype, *N_s_*; carry simultaneously a vaccine serotype and a weak non-vaccine serotype, *B_w_*; or carry simultaneously a vaccine serotype and a strong non-vaccine serotype, *B_s_* (see text). Once vaccinated, individual not carrying vaccine serotypes move to the corresponding states (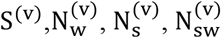) while those carrying vaccine serotypes (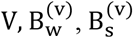) move to the corresponding hyporesponse-associated states (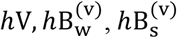) The acquisition rates from the single to multiple serotype carriage states are reduced by competition parameters denoted by *c* with two subscripts; the first denoting the serotype group (*v*,*s and w*, for VT, strong NVT and weak NVT respectively) of the resident serotypes and the second denoting the age-group. The competition parameters have two sets of values, one for age group <6 and another for age group ≥6 years (see Appendix). The age-group specific VT, weak NVT and strong NVT clearance rates are denoted by r_Vi_, r_Nwi_ and r_Nsi_, respectively. In addition to the transitions between the 14 epidemiological states as shown in the Figure, individuals die from any states at age-specific death rates and new individuals are born into the completely susceptible state.

## Appendix Model structure and parameters estimates

### Model structure

A more detailed description of the model and the likelihood function is presented in^1^. The brief description provided in this appendix is to help in the understanding of the notation used, without necessarily referring to^1^. The model is compartmental, age-structured and dynamic. Compartments are defined according to pneumococcal carriage states (Supplementary Figure 2). It has a Susceptible-Infected-Susceptible (SIS) structure for three serotype groups: the PCV10 serotypes, strong NVT and weak NVT.

At any point in time, an unvaccinated individual can be susceptible (non-carrying) in state S; carry a VT, V; carry a weak NVT, N_w_; carry a strong NVT, N_s_; carry simultaneously a weak and strong NVT, N_sw_; carry simultaneously a VT and weak NVT, B_w_; or carry simultaneously a VT and a strong NVT, B_s_. Once vaccinated, the individual moves to one of the corresponding states(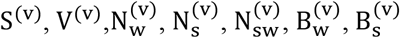). We also fitted a model in which the efficacy of the vaccine on carriage acquisition is reduced due to prevailing carriage at the point of vaccination (hyporesponsiveness) is considered. Under this model, upon vaccination, individuals not carrying vaccine serotypes move to the corresponding states(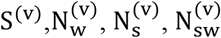) while those carrying vaccine serotypes (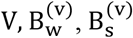) move to the corresponding hyporesponse-related states (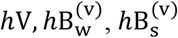) (Supplementary Figure 3).

### Parameterisation

A susceptible unvaccinated individual in age group i becomes colonised with VTs, strong NVTs or weak NVTs at age-group-specific time-dependent rates (forces of infection) denoted by λ(t), λ_Vi_(t), λ_Nsi_(t)and λ_Nwi_(t), respectively. The forces of infection were expressed as functions of the social mixing matrix and age-group specific factors (q_i_) that scale the rate of social contacts into infectious contacts. Due to competition between serotypes in colonising the nasopharynx, the acquisition rate of a secondary serotype is lower than the acquisition rate of that serotype in a completely susceptible individual. Three competition parameters, c_v0_, c_w0_ and c_s0_, represent the fraction by which acquisition rates of secondary serotypes are reduced in <6 year olds infected with VTs, weak NVTs and strong NVTs, respectively. Two competition parameters, c_vw_ = c_v_ = c_w_ and c_s_, were used for individuals aged ≥ 6 years infected with VTs/weak NVTs and strong NVTs, respectively. In the vaccinated compartments the rate of acquisition of VTs are reduced by the vaccine efficacy against carriage acquisition denoted *ε*, or *ε_h_* according to whether the compartment is associated with hyporesponsiveness.

**Table A1.** Estimated parameters of the dynamic transmission models

| Parameter | Estimate (95% Credible Interval) | Estimate accounting for<br>hyporesponsiveness (95%<br>Credible Interval) |
| --- | --- | --- |
| Competition parameters | $c_{s0} = 0.42$ (0.24, 0.62)<br>$c_{w0} = 0.73$ (0.44, 0.97)<br>$c_{v0} = 0.44$ (0.25, 0.70)<br>$c_s = 0.11$ (0.01, 0.40)<br>$c_{vw} = c_v = c_w = 0.70$ (0.30, 0.98) | $c_{s0} = 0.41$ (0.25, 0.59)<br>$c_{w0} = 0.70$ (0.43, 0.97)<br>$c_{v0} = 0.46$ (0.27, 0.70)<br>$c_s = 0.10$ (0.01, 0.30)<br>$c_{vw} = c_v = c_w = 0.66$ (0.24, 0.98) |
| Probability of infection per 100 contacts | $q_1 = 0.14$ (0.11, 0.19)<br>$q_2 = 0.45$ (0.38, 0.55)<br>$q_3 = 0.30$ (0.26, 0.35)<br>$q_4 = 0.08$ (0.06, 0.11)<br>$q_5 = 0.16$ (0.13, 0.19)<br>$q_6 = 0.06$ (0.05, 0.07) | $q_1 = 0.14$ (0.11, 0.19)<br>$q_2 = 0.45$ (0.39, 0.54)<br>$q_3 = 0.30$ (0.26, 0.35)<br>$q_4 = 0.08$ (0.06, 0.10)<br>$q_5 = 0.16$ (0.13, 0.19)<br>$q_6 = 0.06$ (0.05, 0.07) |
| Vaccine efficacy again carriage | $\varepsilon = 0.59$ (0.49, 0.68) | $\varepsilon = 0.58$ (0.47, 0.68) |
| Vaccine efficacy against carriage for VT carriers (hyporesponsiveness) | N/A | $\varepsilon_h = 0.54$ (0.40, 0.68) |

The Metropolis-Hastings algorithm was used to draw samples from the posterior distributions of the parameters. Uniform priors in the range 0-1 were used for competition parameters and the social contact scaling parameters (q_i_). For the vaccine efficacy parameters we used a normal prior centered around 50% with 95% uncertainty interval of 40-60%. 50,000 adaptive MCMC iterations were used. After a burn-in of 25,000 was discarded the remaining stationary samples were thinned to 5000 to estimate the posterior distribution. Convergence was assessed graphically, by observing was no negative or positive trend (zero gradient) in the chain, and by using Geweke diagnostic to check if a chain was stationary. The thinned posterior samples of the parameters were summarised to obtain point estimates (posterior mean) and probability (credibility) intervals. The parameter estimates are shown in Table A1.

